# Macrophage Inflammatory State Influences Susceptibility to Lysosomal Damage

**DOI:** 10.1101/2020.05.29.122390

**Authors:** Amanda O. Wong, Matangi Marthi, Irene A. Owusu, Christiane E. Wobus, Joel A. Swanson

## Abstract

Macrophages possess mechanisms for reinforcing the integrity of their endolysosomal membranes against damage. This property, termed inducible renitence, was previously reported for macrophages stimulated with LPS, peptidoglycan, IFN-γ, or TNF-α. Here, we expanded the macrophage subtypes examined to include populations with well-defined functional roles *in vivo*: classically activated macrophages (CA-Mφ), alternatively activated macrophages (AA-Mφ), and regulatory macrophages (Reg-Mφ). We determined that renitence is a property of CA-Mφ and Reg-Mφ, but not of AA-Mφ. Furthermore, LPS-activated macrophages possess features of both CA-Mφ and Reg-Mφ, based on their cytokine secretion profiles. As the generation of these three classes of renitent macrophages required exposure to LPS, a Toll-like receptor (TLR) ligand, we assessed whether TLR stimulation generally induced renitence. Stimulation of TLRs 2/1, 3, and 4 induced renitence, whereas stimulation of TLRs 7/8 and 9 induced modest levels of lysosomal damage protection. Renitence induced by TLR stimulation required the signaling adaptors MyD88 and TRIF. Surprisingly, the specific signaling adaptor usage requirements for some TLRs differed from those established for canonical TLR signaling. Of note, renitence induced by LPS, a TLR4 ligand, required signaling through TRIF but not MyD88. Consistent with this pattern, the type I IFN response, which is triggered by LPS stimulation through a TRIF-dependent, MyD88-independent pathway, contributed to renitence. A biologically relevant type I IFN trigger in macrophages, murine norovirus-1 (MNV-1) infection, also induced renitence. This work establishes the concept that susceptibility to lysosomal damage within macrophages varies according to inflammatory state and depends on the type I IFN response.

**Summary sentence:** Macrophages of distinct polarization states exhibit varying degrees of susceptibility to lysosomal damage reflecting their functional roles in host defense and immune regulation

## Introduction

Macrophages are professional phagocytes with diverse roles in immunobiology. These range from the clearance of apoptotic bodies to the elimination of pathogenic microbes to the regulation of immune responses. To achieve these diverse functions, macrophages exhibit enormous functional heterogeneity and plasticity [1, 2]. The functional state a macrophage assumes is influenced by the tissue in which it resides and the signals it receives within that environment. Extensive efforts have characterized several functional classes with distinct roles *in vivo*. These include classically activated macrophages (CA-Mφ), alternatively activated macrophages (AA-Mφ), and regulatory macrophages (Reg-Mφ), which have specialized functions in host defense, wound healing, and dampening of immune responses, respectively [1]. These macrophages can be generated *in vitro* through exposure to the same polarizing cytokines that induce their generation *in vivo*. CA-Mφ are generated through stimulation of interferon-γ (IFN-γ) and tumor necrosis factor-α (TNF-α), or with Toll-like receptor (TLR) agonists that induce macrophages to secrete TNF-α. IL-4 and/or IL-13 stimulation induces the generation of AA-Mφ. TLR stimulation coupled with a second signal that reprograms macrophages to adopt an immunosuppressive phenotype, such as that provided by IgG immune complexes, prostaglandin E2 (PGE2), or adenosine (Ado), generates Reg-Mφ [1, 3].

An emerging body of work has begun to clarify the cell biological basis of the phenotypic differences between macrophages of various polarization states. For example, based on studies of human macrophages, the kinetics and extent of phagosome maturation within CA-Mφ and AA-Mφ differ significantly and may underlie differences in their functional roles [4]. Phagosomes within AA-Mφ acidify quickly whereas those in CA-Mφ acidify to a lesser extent, likely reflecting the advantage of promoting rapid clearance of apoptotic bodies in the former and of prolonging exposure of pathogens to phagosome-generated reactive oxygen species (ROS) in the latter [4]. In fact, uptake of a given phagocytic target by macrophages of the inappropriate class can have deleterious consequences. Uptake of apoptotic bodies by CA-Mφ can lead to presentation of self-antigen and development of autoimmune pathologies [5]. AA-Mφ, in contrast to CA-Mφ, are permissive rather than restrictive of many infections [6]. Thus, while the unifying function of macrophages is their ability to perform phagocytosis, the immunological context in which phagocytosis takes place strongly influences the fate of the phagocytic target intracellularly and the cellular consequences for the host.

Here we examined whether the same principle applies for another cellular property of macrophages – namely, their susceptibility to lysosomal damage. Previous work in our lab uncovered a novel macrophage activity called “inducible renitence,” which describes the enhanced ability of macrophages stimulated with lipopolysaccharide (LPS), peptidoglycan, IFN-γ, or TNF-α to resist damage to their phagolysosomes following the phagocytosis of membrane-damaging silica beads [7]. As phagolysosomal damage represents a common threat posed by pathogens, and as the factors found to induce renitence correspond to microbial ligands or host pro-inflammatory cytokines, we reasoned that renitence is a consequence of classical macrophage activation. However, other types of activated macrophages not yet examined might have mechanisms for limiting damage to their lysosomes. Here, we sought to determine the range of macrophages subtypes capable of inducing renitence and to further our understanding of how macrophage cell biological function correlates with inflammatory state.

Of the macrophage activation states examined in previous work, macrophages stimulated overnight with LPS exhibited the most pronounced and consistent protection against lysosomal damage [7]. While LPS-activated macrophages, like CA-Mφ, have been noted for their anti-microbial properties [8, 9], their precise activation state has not been well-defined. That they are generated in the absence of IFN-γ suggests they may not represent canonical CA-Mφ, whose generation *in vitro* occurs under substantially different conditions – namely, several hours of IFN-γ priming followed by overnight stimulation with LPS and IFN-γ [10]. Supporting this idea, TLR stimulation of macrophages in the absence of IFN-γ priming induces the differentiation of macrophages that initially resemble CA-Mφ in terms of their pattern of cytokine secretion, but that over several hours transition to an immunosuppressive state [11]. The regulatory cascade driving this transition has been proposed to serve as an autoregulatory mechanism used by macrophages to prevent the development of hyperinflammatory responses following TLR activation [12]. According to this model, the release of pro-inflammatory cytokines following TLR stimulation is accompanied by the release of low levels of adenosine triphosphate (ATP), which eventually is converted into adenosine (Ado), a signal that promotes the generation of immunosuppressive Reg-Mφ [12, 13]. Ado acts as a ligand for the adenosine 2b receptor (A2bR), whose activation triggers downstream signaling leading to the expression of Reg-Mφ genes [13]. This reprogramming step is avoided in macrophages primed with IFN-γ, as IFN-γ inhibits the expression of A2bR, such that cells become unresponsive to Ado [11].

Based on the above model, we predicted that macrophages observed previously to undergo renitence (namely, LPS-activated macrophages stimulated in the absence of IFN-γ priming) may harbor characteristics of Reg-Mφ, and, as a corollary, that Reg-Mφ likely exhibit renitence. We sought to test these predictions within the context of two broader aims: (1) to expand our understanding of the immunological contexts in which renitence acts, and (2) to define and compare the inflammatory states of macrophages that do or do not induce renitence.

We report that only a subset of macrophage subtypes induced renitence. These included CA-Mφ, Reg-Mφ, and macrophages treated with ligands of TLRs 2/1, 3, and 4. Macrophages that did not induce or only modestly induced renitence included AA-Mφ and macrophages treated with ligands of TLRs 7/8 and 9. Building upon these observations, we examined the dependence of TLR signaling for renitence, the contribution of the type I IFN response to renitence, and the potential for viral infection to induce renitence. Together, this work expands our understanding of the conditions that induce renitence and supports the concept that the polarization state of macrophages affects their susceptibility to lysosomal damage.

## Materials and Methods

### Mice and macrophage isolation

C57BL/6J mice were purchased from The Jackson Laboratory (Bar Harbor, ME). *Myd88*^-/-^, *Trif*^-/-^, and *Myd88/Trif*^-/-^ mice were provided by Gabriel Nuñez (University of Michigan). All mice were maintained under specific pathogen-free conditions at the University of Michigan. Bone marrow cells isolated from the femurs and tibia of mice were differentiated into bone marrow-derived macrophages (BMM) through culture for 6-8 days in DMEM containing 10% fetal bovine serum (FBS) and 50 ng/ml recombinant M-CSF, as previously described [7]. Femurs and tibia from *Ifnar1*^-/-^ mice on a C57BL/6J background were provided by Megan Baldridge (Washington University in St. Louis, MO, USA). For all experiments using cells from these mice, *Ifnar1*^-/-^ and WT BMM were differentiated through culture for 6 days in L929-conditioned DMEM containing 20% FBS, 30% L9 supernatant, 1% L-Glutamine, 1% Sodium Pyruvate, 0.1% beta-mercaptoethanol and 1% Penicillin/Streptomycin, as described in [14].

### Cell culture and stimulation

CA-Mφ were generated by priming BMM with 150 U/ml IFN-γ (R&D Systems, Minneapolis, MN) for 6 h, and then stimulating cells overnight with 150 U/ml IFN-γ and 100 ng/ml LPS (from *Salmonella typhimurium*; no. 225; List Biological Laboratories, Campbell, CA). AA-Mφ were generated by stimulating macrophages overnight with 20 ng/ml IL-4 (R&D Systems, Minneapolis, MN). Reg-Mφ were generated by stimulating macrophages overnight with 100 ng/ml LPS and either 200 nM PGE2 (Cayman Chemical, Ann Arbor, MI) or 200 μM adenosine (Sigma-Aldrich, St. Louis, MO). Studies of macrophages stimulated overnight with TLR agonists were performed with the following reagents: Pam3CSK4 (100 ng/ml); poly(I:C) (10 μg/ml); ultrapure flagellin from Salmonella typhimurium (FLA-ST; 100 ng/ml); R848 (100 ng/ml), ODN 1826 (1 μM). All TLR agonists were purchased from Invivogen (San Diego, CA) except for poly(I:C), which was purchased from Tocris (Bristol, United Kingdom).

For experiments in which both RNA isolation and cytokine analyses were performed, 6 x 10^6^ cells were plated onto 60-mm dishes (ThermoFisher, Waltham, MA). For experiments in which cytokine analyses were performed and RNA was not isolated, 1 x 10^5^ cells were plated onto 24-well plates (ThermoFisher). For assays of lysosomal damage, 8 x 10^4^ cells were plated onto 35-mm dishes with attached 14-mm coverglass (MatTek Corporation, Ashland, MA).

### Gene expression analysis

RNA was isolated using a Qiagen RNeasy Mini Kit (74104; Venlo, Netherlands) and converted into cDNA using MMLV-Reverse Transcriptase from ThermoFisher (28025013). Quantitative PCR (qPCR) analysis was performed using an Applied Biosystems 7500 Fast Real-Time PCR system (ThermoFisher) and Brilliant II SYBR Green Master Mix (600830; Agilent, Santa Clara, CA). Primer pairs used for amplification of specific gene products are as follows: *Il-12p40* F, AAGACGTTTATGTTGTAGAGGTGGAC; *Il-12p40* R, ACTGGCCAGGATCTAGAAACTCTTT; *Il-10* F, GACTTTAAGGGTTACTTGGGTTGC; *Il-10* R, TCTTATTTTCACAGGGGAGAAATCG; *Relm-α* F, AATCCAGCTAACTATCCCTCCA; *Relm-α* R, CAGTAGCAGTCATCCCAGCA; *Gapdh* F, AAGGTCGGTGTGAACGGATTT; *Gapdh* R, AATTTGCCGTGAGTGGAGTCATAC. Primers are listed 5’ to 3’ and were previously published in [3]. Relative expression levels were calculated using the ΔΔCT method, using *Gapdh* as the reference gene for normalization [15].

### Cytokine measurements

Mouse IL-12p40, mouse TNF-α, and mouse IL-10 cytokine concentrations were determined using ELISA DuoSet kits (R&D Systems, Minneapolis, MN).

### Particle preparation

3 μm diameter silica dioxide microspheres were purchased from Microspheres-Nanospheres, a subsidiary of Corpuscular Inc (Cold Spring, NY). To clean the particles of debris, microspheres were acid-washed (AW) overnight in 1N HCl, then rinsed several times with Milli-Q-filtered water.

### Measurement of lysosomal damage by ratiometric imaging

BMM were plated onto glass-bottom MatTek dishes in RPMI 1640 containing 10% FBS, 1% GlutaMAX supplement, and 10 U/ml penicillin-streptomycin. Damage to macrophage lysosomes was measured using an assay for ratiometric measurement of pH. To label macrophage lysosomes, BMM were incubated overnight with 150 μg/ml fluorescein dextran, average molecular weight 3 kDa (Fdx; ThermoFisher). During this overnight pulse, cells also were treated with the appropriate polarizing cytokines for generating various macrophage subtypes. The next day, cells were rinsed in Ringer’s buffer (155 mM NaCl, 5 mM KCl, 2 mM CaCl2, 1 mM MgCl2, 2 mM NaH2PO4, 10 mM HEPES, and 10 mM glucose) and returned to unlabeled media for at least 3 hours before the start of imaging. Lysosomal damage was induced by feeding BMM 3 μm diameter AW beads in RPMI lacking serum for 60 min. AW beads were added at a concentration empirically determined to result in uptake of on average 3 to 4 beads per cell by both resting and LPS-activated BMM. All analyses of damage were performed on cells that had internalized 3 to 7 beads per cell.

To monitor dye release, BMM containing Fdx-labelled lysosomes were imaged by fluorescence microscopy after 60 min incubation in the presence or absence of AW beads. For each field of cells imaged, three images were acquired: a phase-contrast image, which allowed enumeration of bead number per cell, and two fluorescent images, captured using a single emission filter centered at 535 nm and two excitation (exc.) filters, centered at 440 nm or 490 nm, the pH-insensitive and pH-sensitive wavelengths, respectively, for fluorescein. Taking the ratio of 535 nm fluorescence intensities captured at exc. 490 and exc. 440 yielded pH information for each pixel in the image. A calibration curve was generated by measuring 490 nm/440 nm excitation ratios of Fdx in BMM exposed to the ionophores nigericin and valinomycin (10 μM) in fixed pH clamping buffers [7]. Release of dye from lysosomes was quantified as the percent of pixels residing in cellular regions whose pH was greater than 5.5. This measurement of percent Fdx release was made on a per-cell basis and reported as the average percent Fdx release for each condition. Image acquisition and analysis were performed using Metamorph software (Molecular Devices, Sunnyvale, CA) as described in [16].

### Virus stock

The plaque-purified MNV-1 clone (GV/MNV1/2002/USA) MNV-1.CW3 [17] (referred to here as MNV-1) was used at passage 6 in all experiments. Virus-containing cell lysate was titered by plaque assay on RAW 264.7 cells as described in [18].

### Viral infection and measurement of viral titers

BMM were seeded overnight on MatTek dishes for lysosomal damage assays and in parallel in 24-well plates for measurement of viral titers. Cells were infected with MNV-1 stock at multiplicities of infection (MOI) of 0.05, 0.5 and 5, then kept on ice for 1 hour with gentle shaking. Inoculum was removed and cells were washed twice with cold DPBS with calcium and magnesium and replaced with RPMI 1640 containing 10% FBS, 1% GlutaMAX supplement, and 10 U/ml penicillin-streptomycin. After 18 h of infection, cells were subjected to assays of lysosomal damage, as described above. BMM infected in parallel were replaced in DMEM containing 10% FBS, 10% L9 supernatant, 1% L-Glutamine, 1% non-essential amino acids and 1% HEPES buffer solution and freeze-thawed twice. MNV-1 titer was determined using plaque assays on RAW 264.7 cells as described in [19].

### Statistical methods

Statistical analysis was performed using GraphPad Prism software (GraphPad Software Inc; La Jolla, CA). For gene expression and cytokine secretion analyses, statistical significance relative to unstimulated cells was determined using an unpaired, two-tailed, nonparametric t-test (Mann-Whitney). For analyses of lysosomal damage, the average percent Fdx release was calculated from pooled individual cell data for each condition. Typically, two to five independent experiments were performed per condition resulting in a n of > 100 cells per condition. The same data sets were used to generate histograms displaying the frequency distributions of lysosomal damage for each condition. Average percent Fdx release values between groups were compared using two-way ANOVA with Tukey’s multiple comparisons. Statistical difference between each pair of frequency distributions was assessed using an unpaired, two-tailed, non-parametric t-test (Kolmorogov-Smirnov).

## Results

### Generation and characterization of variously activated macrophages

CA-Mφ, AA-Mφ, and Reg-Mφ were generated and assessed for their ability to undergo renitence. To generate macrophages of each subtype, we exposed murine bone marrow-derived macrophages (BMM) in culture to the appropriate polarizing cytokines. By gene expression and cytokine secretion analysis, we confirmed that these treatments successfully generated macrophages of the expected subtypes [10]. LPS and IFN-γ treatment of macrophages first primed for 6h with IFN-γ generated CA-Mφ producing high levels of IL-12p40 and TNF-α, and low levels of IL-10 (Fig. 1A and B). Macrophages treated with LPS in combination with either prostaglandin E2 (PGE2) or adenosine (Ado) generated Reg-Mφ producing low levels of IL-12p40 and TNF-α, and high levels of IL-10. Finally, IL-4 stimulation of macrophages yielded AA-Mφ producing low levels of IL-12p40 and IL-10, but expressing high levels of *Relm-α* (Fig. 1A). As expected, this gene was not abundantly expressed in either CA-Mφ or Reg-Mφ.

**Figure 1.**
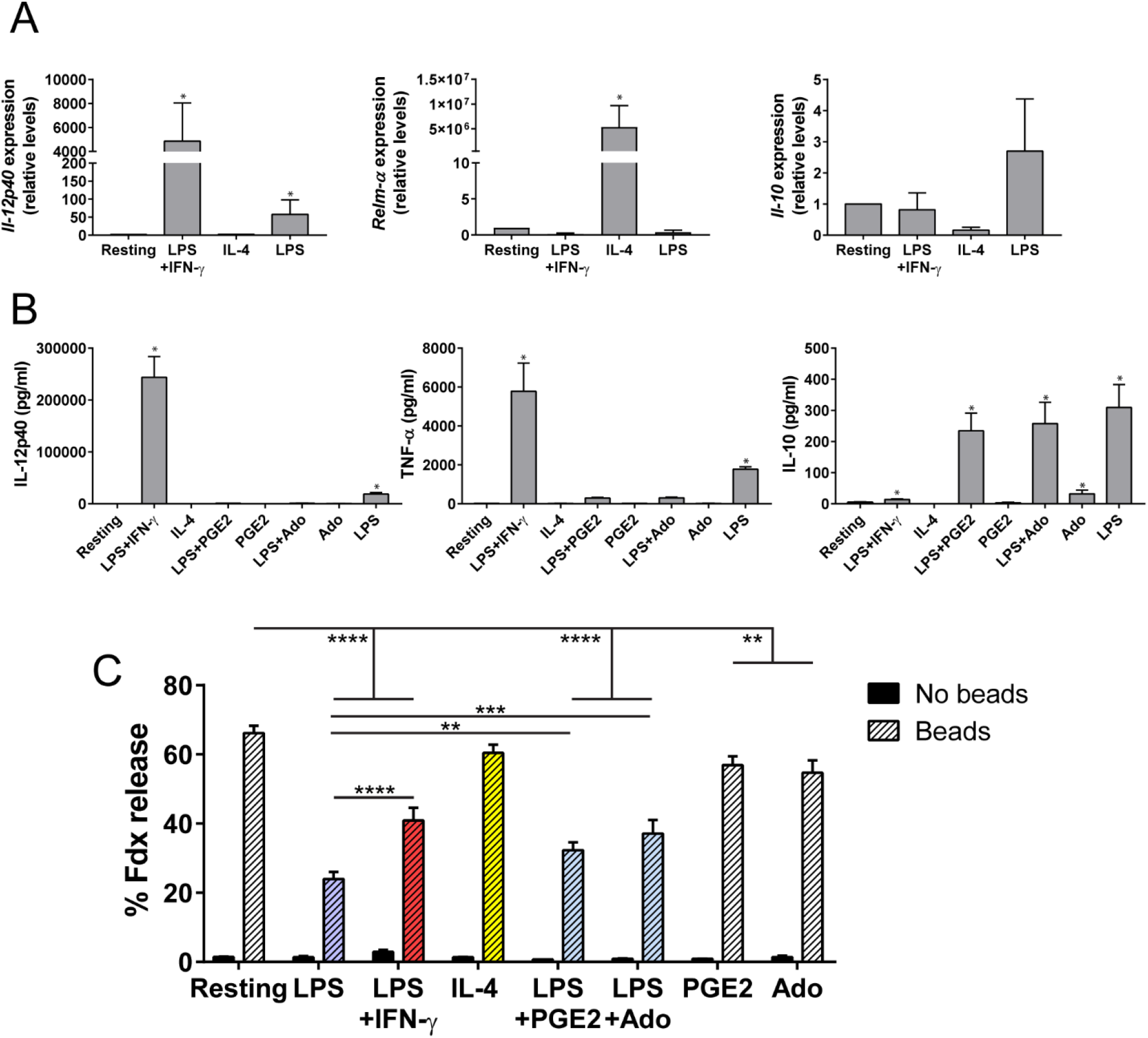
Renitence is a property of classically activated and regulatory macrophages. (A) BMM were treated overnight with LPS and IFN-γ (after initial 6 h IFN-γ priming), IL-4, or LPS alone, or left unstimulated (Resting). For each condition, mRNA expression of *Il-12p40*, *Relm-α*, and *Il-10* relative to levels expressed in resting BMM was determined by qPCR. Bars represent mean ± SEM calculated from two (*Il-10*) or three (*IL-12p40*, *Relm-α*) independent experiments. Statistical significance relative to expression levels in resting BMM is indicated. *p ≤ 0.05. (B) BMM were subjected to the following treatments for generating classically activated (CA-Mφ), alternatively activated (AA-Mφ), or regulatory macrophages (Reg-Mφ): 6 h IFN-γ priming followed by overnight stimulation with LPS and IFN-γ to generate CA-Mφ; overnight stimulation with IL-4 to generate AA-Mφ; overnight stimulation with LPS in combination with prostaglandin E2 (PGE2) or adenosine (Ado) to generate Reg-Mφ. As controls, macrophages were left unstimulated (Resting) or treated overnight with LPS, PGE2, or Ado alone. Levels of IL-12p40, TNF-α, and IL-10 in cell supernatants were measured by ELISA. Each bar represents mean ± SEM of at least three independent experiments. Statistical significance relative to levels of cytokine secretion in resting BMM is shown. *p ≤ 0.05. (C) BMM were subjected to the indicated treatments for generating CA-Mφ (red), AA-Mφ (yellow), Reg-Mφ (blue), or control macrophages, which included resting macrophages and BMM singly treated with LPS (purple), PGE2, or Ado. Concurrent with receiving the indicated treatments, BMM in each condition were pulsed overnight with fluorescein dextran (Fdx) to label the endolysosomal network. The next day, cells were chased in unlabeled medium for 3 h to allow Fdx movement into lysosomes. To initiate membrane damage, cells were incubated with acid-washed (AW) beads for 60 min or received no bead challenge as controls. Ratiometric imaging was performed to measure the extent of Fdx release from lysosomes. Bars represent the average percent Fdx release ± SEM per condition. In the groups of cells receiving beads, analysis was restricted to cells internalizing three to seven beads. Shown are pooled cell data from two to four independent experiments (n > 97 cells per condition). **p ≤ 0.01, ****p ≤ 0.0001.

While macrophages stimulated overnight with LPS are commonly regarded as CA-Mφ, their activation status has not been precisely defined. We determined that macrophages stimulated overnight with LPS exhibit features of both CA-Mφ and Reg-Mφ. In addition to producing high levels of IL-12p40 and TNF-α, LPS-treated macrophages also produced high levels of IL-10, to a similar extent to that produced by Reg-Mφ (Fig. 1B). Placed on the spectrum of macrophage activation, macrophages stimulated overnight with LPS assume an intermediate phenotype between that of CA-Mφ and Reg-Mφ.

### Classically activated and regulatory macrophages exhibit renitence

We next assessed the ability of macrophages of each functional class to undergo renitence. We previously defined renitence as an inducible activity within macrophages that confers protection against lysosomal damage [7]. Our methods for inducing and measuring lysosomal damage allow for the sensitive detection of leakage of lysosomal contents in cells subjected to comparable levels of injury. Damage induction was achieved through challenging cells with silica beads, which upon phagocytosis have the potential to damage endolysosomal membranes. Based on previous work, silica bead challenge mimics the extent and timing of phagolysosomal damage caused by pathogens [16] but avoids the confounding effects introduced by study of microbial pathogens. Measurements of lysosomal damage were performed using a live-cell, ratiometric fluorescence microscopy-based assay [20]. Prior to the onset of damage induction, BMM lysosomes were pulse-chase labeled with the pH-sensitive dye, fluorescein dextran (Fdx, 3 kDa average molecular weight). Cells were then subjected to lysosomal damage through challenge with acid-washed, 3 μm diameter silica beads (AW beads). After a 60 min incubation with the AW beads, ratiometric fluorescence imaging was performed to determine the proportion of the dye that was released from lysosomes into the cytoplasm within individual cells. To ensure comparison of cells receiving similar levels of injury, analysis was restricted to cells containing three to seven beads per cell. Damage to lysosomes was quantified on a per-cell basis and reported as the average percent release of Fdx for all cells analyzed in a given condition. For a more granular analysis, histograms depicting the distribution of damage based on the individual cell data for each population were compared.

As previously demonstrated [7, 16], resting macrophages challenged with AW beads experienced considerable damage that was significantly reduced in cells pre-treated with LPS (Fig 1C). CA-Mφ, as well as Reg-Mφ generated through two different treatments (LPS+PGE2; LPS+Ado), showed protection from AW bead-mediated damage, but to a lesser degree than that seen in LPS-treated macrophages. The protective effect seen in Reg-Mφ was dependent on the presence of LPS, as single treatment with either PGE2 or Ado conferred only modest protection. AA-Mφ, unlike CA-Mφ or Reg-Mφ, did not exhibit protection from AW bead-mediated damage, experiencing similar levels of damage as that seen in resting macrophages. Together, these results demonstrate that renitence is a property of CA-Mφ and Reg-Mφ, but not of AA-Mφ.

Plotting the individual cell data from Figure 1C into histograms that depict the relative frequency of cells experiencing a given level of lysosomal damage reveals that cells experience damage heterogeneously with frequency distributions that are skewed toward high damage (right-most bars in Supplementary Figures 1-5) or low damage (left-most bars). A similar observation was made through comparing the frequency distributions of % Fdx release values in resting and LPS-treated BMM after various time points of AW bead incubation [16]. Through that analysis, we discovered that most cells in either population experienced either complete (90-100%) or negligible (0-10%) Fdx release. Resting BMM have a higher average level of damage, first detected after 30 min silica bead challenge, that is accounted for by a large proportion of cells experiencing very high levels of damage and a small proportion of cells experiencing very low levels of damage. Conversely, over time LPS-treated BMM maintained a high percentage of cells experiencing very low levels of damage and a low percentage of cells experiencing very high levels of damage. From this work we concluded that LPS-treated macrophages limit lysosomal damage within a population by preventing a protected subpopulation of cells from undergoing damage [16].

We plotted the frequency distribution of damage levels for each polarization state examined in Figure 1C against that for resting BMM. By visualizing the data in this way, we found that all polarization states determined to exhibit renitence based on comparison of average % Fdx release values in Figure 1C (LPS, LPS+IFN-γ, LPS+PGE2, LPS+Ado) exhibited frequency distributions of similar shape and spread (Suppl Fig 1A, B, D, E). In each of these conditions, the group of cells with the highest relative frequency were those undergoing 0-10% damage. As the majority of cells fell into this category, the frequency distributions for these conditions were skewed to the left. In contrast, conditions found not to exhibit renitence in Figure 1C (IL-4, PGE2, Ado) had frequency distributions that visually resembled that for resting BMM and were skewed to the right (Suppl Fig 1C, F, and G). Thus, polarization states that exhibit renitence seem to restrict damage by preserving lysosomal integrity within a predominant subpopulation of cells.

The pattern of protected versus unprotected subsets suggests that renitence is an activity characteristic of macrophages specialized in host defense either in their present state (CA-Mφ) or recent past (Reg-Mφ). Indeed, CA-Mφ and Reg-Mφ share a common requirement for their generation: exposure to a microbial ligand – namely, LPS. As LPS activates downstream signaling in macrophages through stimulating TLR4, we next asked whether stimulation of other TLRs also confers protection against lysosomal damage in macrophages.

### TLR stimulation by a subset of agonists induces renitence

We previously observed that stimulating cells with peptidoglycan, a TLR2 agonist, induced renitence to a similar degree as that induced by LPS [7], suggesting that TLR stimulation induces renitence. To address this possibility, we exposed BMM to a panel of TLR agonists, and assessed the ability of each agonist to protect macrophage lysosomes from AW bead-mediated injury. Only a subset of the tested agonists induced renitence. These included the synthetic triacylated lipopeptide Pam3CSK4, a bacterial ligand that activates TLR2/1; poly(I:C), an analog of dsRNA, which activates TLR3; and LPS, a component of the cell wall of gram-negative bacteria and canonical TLR4 ligand (Fig. 2A). R848 (Resiquimod), an anti-viral compound that activates TLR7/8, and ODN 1826, a synthetic oligonucleotide containing unmethylated CpG motifs, which activates TLR9, induced much more modest protection against lysosomal damage. Flagellin purified from *S. typhimurium* (FLA-ST), an agonist of TLR5, appeared to offer no protection from lysosomal damage. However, based on cytokine secretion analysis, FLA-ST stimulation did not induce secretion of TNF-α, an expected secreted product following TLR stimulation (Fig. 2B). In contrast, all other TLR agonists tested induced robust secretion of TNF-α and variable levels of secretion of other probed cytokines (IL-12p40 and IL-10). FLA-ST thus seems to have failed to activate its cognate receptor, TLR5, in our experiments. The inability of FLA-ST to activate BMM has been noted previously by others and is attributed to low levels of TLR5 expression in mouse BMM [21].

**Figure 2.**
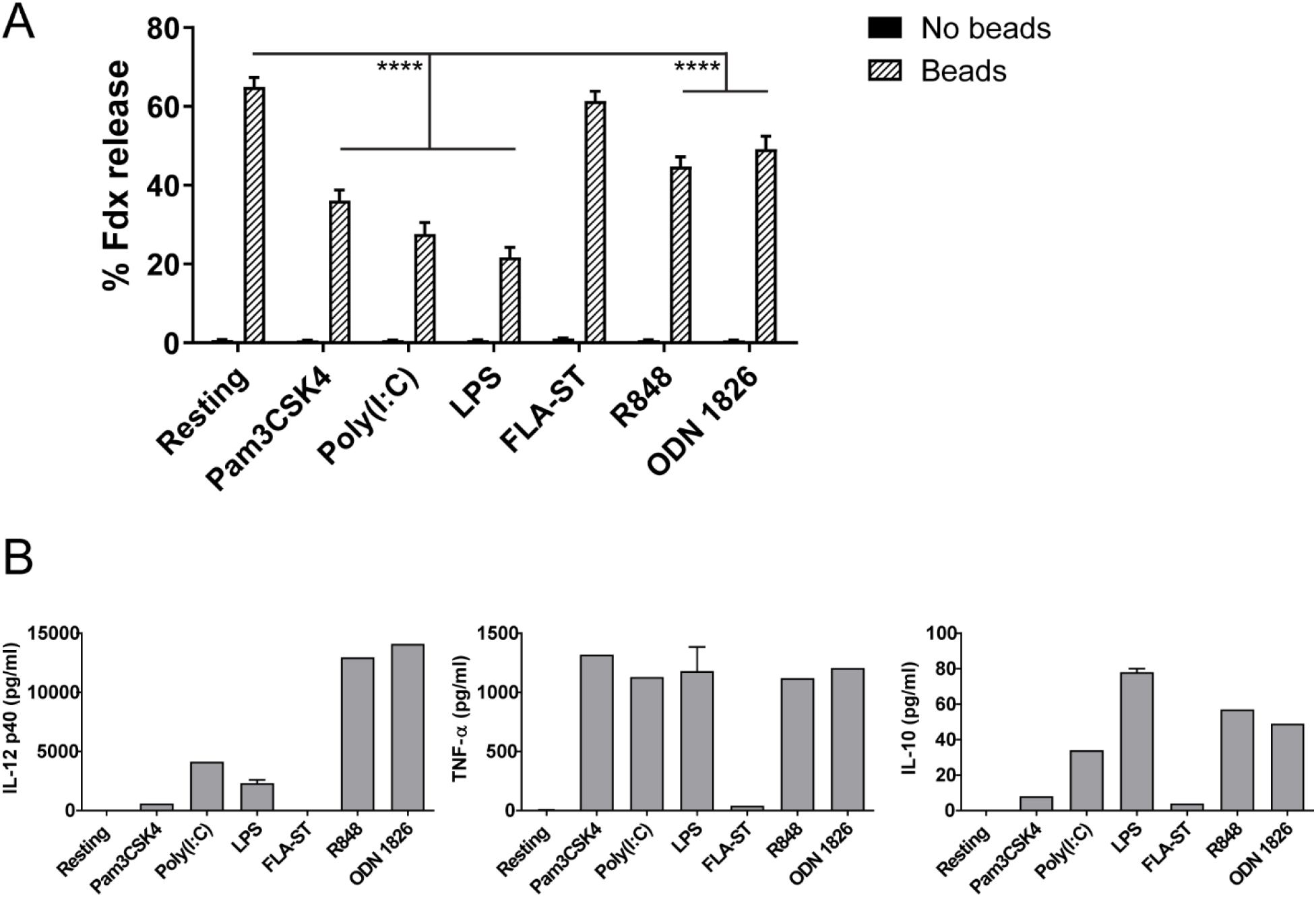
A subset of TLR agonists induces renitence. (A) BMM were pulsed overnight with Fdx while undergoing stimulation with the indicated TLR agonists or left untreated. The next day, cells were chased for 3 h, incubated with AW beads or no beads for 60 min, and imaged to quantify the extent of Fdx release. Bars represent the average percent Fdx release ± SEM per condition. In bead-positive conditions, cells containing three to seven beads were analyzed. Shown are pooled cell data from two or three independent experiments (n > 110 cells per condition). ****p ≤ 0.0001. (B) BMM were stimulated overnight with the indicated TLR agonists or left untreated. Levels of IL-12p40, TNF-α, and IL-10 in cell supernatants were measured by ELISA. Data shown is from one experiment.

Examination of the frequency distributions for each condition in Figure 2 finds that the set of TLR agonists that induced robust renitence (Pam3CSK4, poly(I:C), and LPS) restricted damage through maintaining a large proportion of cells experiencing 0-10% damage (Suppl Fig 2A, B, C). Stimulation with TLR agonists conferring more modest protection (R848 and ODN 1826) failed to maintain as high of a proportion of cells experiencing very low levels of damage, but led to a substantially lower proportion of cells experiencing very high levels of damage compared to that seen in resting BMM (Suppl Fig 2E, F). The frequency distribution for BMM stimulated with FLA-ST, which did not induce renitence, resembled that for resting BMM (Suppl Fig 2D).

Thus, activation of TLRs 2/1, 3, and 4 induced renitence, whereas activation of TLRs 7/8 and 9 induced a modest level of protection from lysosomal damage. The TLRs belonging to either group differ in terms of which signaling adaptors they recruit, which intracellular compartment they signal from, and which class of ligands they recognize. As such, the pattern revealed does not immediately suggest a mechanistic basis for renitence that relates to any of these variables.

We considered whether the differential ability of TLR agonists to induce renitence might reflect differences in the inflammatory state produced by stimulation with the agonists. That is, does the set of agonists capable of inducing renitence generate macrophages of a different activation state than those agonists that do not induce renitence? By examining cytokine secretion levels in macrophages stimulated with each of the TLR agonists, we determined that all TLR agonists tested (with the exception of FLA-ST) induced similar levels of TNF-α secretion regardless of their ability to induce renitence (Fig 2B). The level of IL-10 secretion induced by the panel of agonists was more variable, but likewise did not correlate with either the ability or inability to induce renitence. Levels of IL-12p40 secretion, however, widely differed between the two groups. The agonists with less renitence-inducing potential (R848 and ODN 1826) induced markedly higher production of IL-12p40 than the set of agonists capable of inducing renitence (Pam3CSK4, Poly(I:C), and LPS) (Fig 2B). These results suggest that IL-12p40 secretion correlates inversely with renitence.

### Renitence induced by TLR ligands requires intact TLR signaling

To determine whether the induction of renitence by TLR ligands depends on the canonical pathways of TLR signaling, we measured lysosomal damage following 60 min AW bead incubation in C57BL/6J (WT) and *Myd88* and *Trif*-deficient (*Myd88/Trif^-/-^*) BMM stimulated with the same panel of TLR agonists used in Figure 2A, excluding FLA-ST. As MyD88 (Myeloid differentiation primary response 88) and TRIF (TIR-domain-containing adaptor-inducing interferon-β) are the major signaling adaptors responsible for propagating signaling downstream of TLR activation, macrophages deficient in these two adaptors lack functional TLR signaling. In WT BMM, stimulation with the panel of TLR agonists induced the same pattern of protection as seen in Figure 2A, except that ODN 1826 stimulation of WT BMM did not confer a statistically significant reduction in lysosomal damage over that seen in resting BMM (Fig 3A). Impairment of TLR signaling in *Myd88/Trif^-/-^* BMM eliminated renitence by all agonists tested except for ODN 1826. Consistent with these results, the frequency distributions for each condition reveal a shift toward values of high damage associated with loss of MyD88 and TRIF for every condition except for *Myd88/Trif^-/-^* BMM stimulated with ODN 1826 (Suppl Fig 3A-F). The finding of no exacerbation of damage in ODN 1826-stimulated *Myd88/Trif*^-/-^ BMM suggests neither signaling adaptor contributes to lysosomal damage responses in ODN 1826-stimulated macrophages, whose susceptibility to lysosomal damage in WT BMM is similar to or slightly reduced compared to that seen in resting macrophages (p ≤ 0.0001 in Fig 2A, p ≤ 0.01 in Fig 3C, no significant difference in Fig 3A and 3B). Interestingly, loss of these signaling adaptors exacerbated damage even in resting BMM, suggesting that some aspect of signaling mediated by MyD88 and TRIF functions to protect non-activated macrophages from lysosomal damage. Together, these results suggest that renitence stimulated by TLR agonists other than ODN 1826 requires functional TLR signaling.

**Figure 3.**
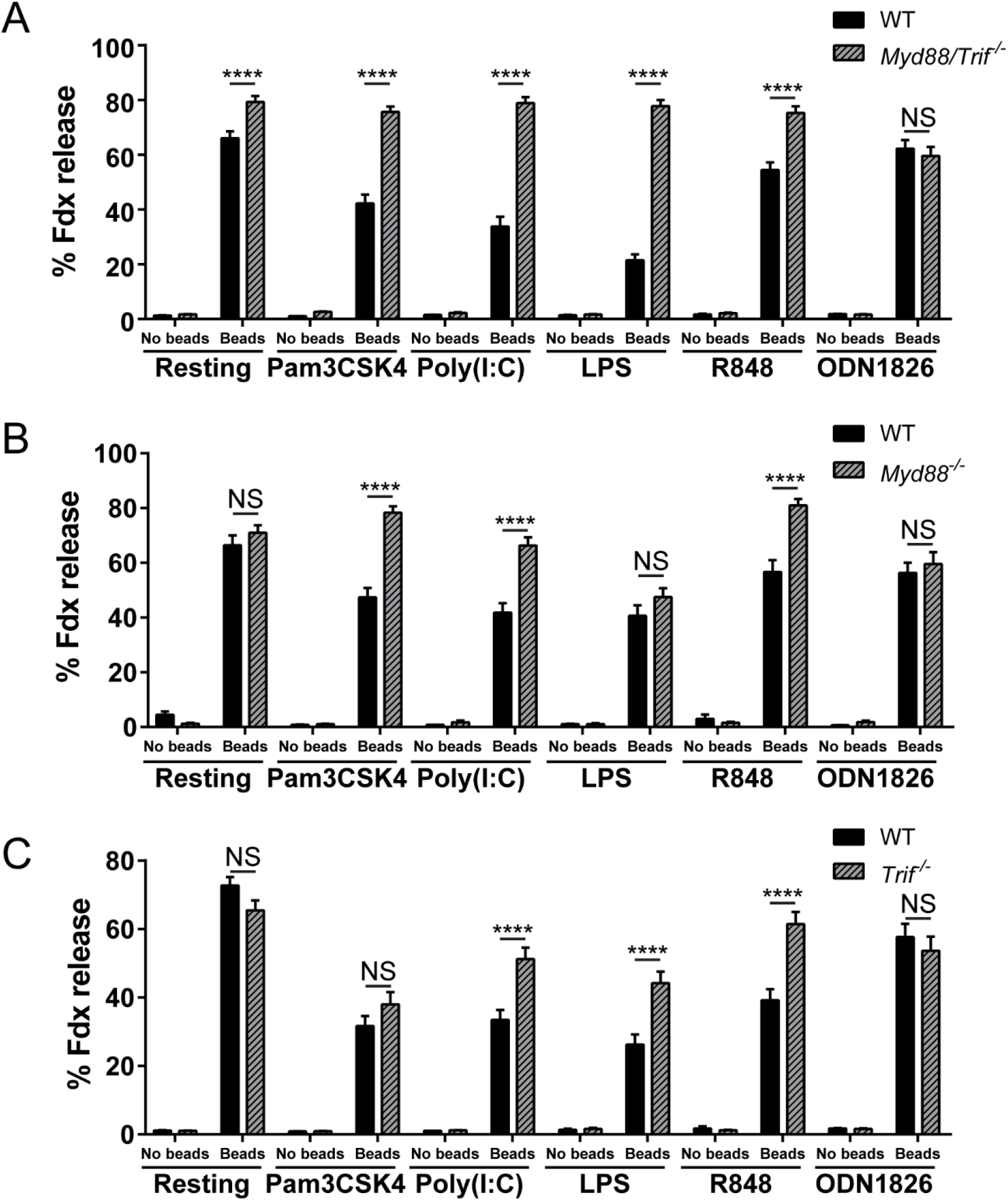
Renitence requires intact TLR signaling, with different adaptor requirements than those for NF-κB signaling. (A) BMM were isolated from C57BL/6J (WT) mice and mice deficient in *Myd88* and *Trif* (*Myd88/Trif*^-/-^). Both groups of BMM were treated overnight with the indicated TLR agonists concurrent with pulse-chase labeling of lysosomes with Fdx. BMM were then incubated with AW beads or no beads for 60 min and assayed for lysosomal damage. Bars represent the average percent Fdx release ± SEM per condition for pooled cell data from two independent experiments. In bead-positive conditions, cells containing three to seven beads were analyzed (n > 107 cells per condition). NS: no significant difference, \*\*\*\**p* ≤ 0.0001. (B) WT and *Myd88*^-/-^ BMM were stimulated with the indicated TLR agonists while subjected to pulse-chase labeling of lysosomes with Fdx. BMM were then incubated with AW beads or no beads for 60 min and assayed for lysosomal damage. Bars represent the average percent Fdx release ± SEM per condition for pooled cell data from two independent experiments. In bead-positive conditions, cells containing three to seven beads were analyzed (n > 63 cells per condition). NS: no significant difference, \*\*\*\**p* ≤ 0.0001. (C) WT and *Trif*^-/-^ BMM were stimulated with the indicated TLR agonists while subjected to pulse-chase labeling of lysosomes with Fdx. BMM were then incubated with AW beads or no beads for 60 min and assayed for lysosomal damage. Bars represent the average percent Fdx release ± SEM per condition for pooled cell data from two independent experiments. In bead-positive conditions, cells containing three to seven beads were analyzed (n > 85 cells per condition). NS: no significant difference, \*\*\*\**p* ≤ 0.0001.

### TLR signaling adaptor requirements for renitence differ from those for NF-κB activation

To dissect which signaling adaptor(s) (i.e. MyD88 and/or TRIF) contribute to renitence induced by each TLR, we measured lysosomal damage in BMM with single gene deficiencies in either *Myd88* or *Trif* (*Myd88*^-/-^ BMM or *Trif*^-/-^ BMM, respectively). We predicted that the pattern of adaptor usage downstream of TLR signaling for renitence would parallel the established patterns of involvement of MyD88 vs. TRIF in TLR signaling leading to NF-κB activation: namely, that TLRs 2/1, 7/8, and 9 signal through MyD88 only, TLR3 signals through TRIF only, and TLR4 signals through both MyD88 and TRIF [22]. Comparing the effect of each individual knockout on renitence induced by each TLR agonist revealed the following adaptor usage requirements for renitence. TLR2/1 stimulation by Pam3CSK4 induced renitence through signaling that requires MyD88 but not TRIF, as renitence was eliminated in *Myd88*^-/-^ BMM (Fig 3B) but unaffected in *Trif*^-/-^ BMM (Fig 3C). In contrast, renitence induced by LPS stimulation of TLR4, whose activation at the cell surface recruits MyD88 as a signaling adaptor and whose activation within endosomes recruits TRIF [23], depended on signaling through TRIF but not MyD88. This result suggests that endosomal but not cell-surface TLR4 signaling is necessary for LPS-induced renitence. Although renitence induced through TLR4 requires TRIF, TRIF deficiency led only to a partial abrogation of protection, suggesting that other pathways downstream of TLR4 signaling contribute to renitence.

While the results described so far are consistent with the known adaptor usage patterns for TLR signaling leading to NF-κB activation, those we obtained for the other TLR agonists tested did not segregate according to these established patterns. Specifically, renitence induced by stimulation of TLRs 3 or 7/8 (receptors for poly(I:C) and R848 respectively) required both MyD88 and TRIF even though canonically TLR3 signals only through TRIF and TLR7/8 signals only through MyD88 (Fig 3B and C). Moreover, deficiency of either MyD88 or TRIF completely abrogated TLR7/8-induced renitence by R848, but only partially abrogated TLR3-induced renitence by poly(I:C). The results obtained for TLR3-induced renitence were therefore unexpected in two ways – first for its requirement for MyD88, which is not involved in canonical TLR3 signaling, and second for the residual renitence observed in BMM deficient in TRIF, whose deficiency based on canonical TLR signaling would be expected to abrogate downstream effector functions. Similarly, how TRIF deficiency leads to the abrogation of renitence induced by TLR7/8 activation when canonical signaling downstream of the receptor does not require TRIF is unclear. The frequency distributions for these conditions show a shift in the distribution toward values of high damage in *Trif*^-/-^ BMM stimulated with R848 (Suppl Fig 3Q) but a more limited shift in *Trif*^-/-^ BMM stimulated with poly(I:C) (Suppl Fig 3O) or LPS (Suppl Fig 3P). In the latter conditions, loss of TRIF led to a reduction in the proportion of cells experiencing very low levels of damage, and an even distribution of damage across all other levels of % Fdx release. Thus, TRIF signaling in poly(I:C) and LPS-stimulated BMM contributes toward the maintenance of a large sub-population of macrophages undergoing low levels of damage but does not seem to affect the proportion of cells undergoing high levels of damage. Finally, lysosomal damage in BMM undergoing TLR9 activation by ODN 1826, was unaffected by knockout of either MyD88 or TRIF. Consistent with this observation, the frequency distributions for ODN 1826-stimulated BMM were unaffected by cell genotype (Suppl Fig 3F, L, R). Thus, the response to lysosomal damage in ODN 1826-stimulated macrophages is independent of signaling through MyD88 or TRIF. Together, these studies demonstrate that the requirement for MyD88 or TRIF for renitence varies depending on the TLR stimulated, and that the pattern of receptor usage necessary for renitence deviates for some TLRs (namely TLRs 3 and 7/8) from the pattern expected for NF-κB activation.

### The type I IFN response contributes to renitence induced by agonists of TLR4 and TLR3

Of the TLR agonists examined here, the two that conferred the greatest protective effect of renitence were LPS and poly(I:C), agonists of TLRs 4 and 3, respectively. As shown in Figure 3C, renitence induced through stimulation of each of these receptors requires the adaptor molecule TRIF. This observation prompted us to consider the contribution of TRIF-dependent pathways to renitence. One such pathway is the type I interferon (IFN) response, originally discovered for its role in anti-viral immunity but more recently also found to be activated by and contribute to immunity against bacteria, parasites, and fungi [24]. Stimulation of either TLR4 or TLR3 induces type I IFN production in many cell types, including macrophages, in a TRIF-dependent, MyD88-independent manner [23, 25]. Stimulation of the other TLRs induces type I IFN production only in select cell types, including plasmacytoid dendritic cells and conventional dendritic cells, and occurs in a MyD88-dependent fashion [25].

To assess the effect of the type I IFN response on renitence induced by stimulation of TLR4 or TLR3, we compared the extent of lysosomal damage in LPS and poly(I:C)-treated WT BMM and *Ifnar1*^-/-^ BMM, which cannot respond to secreted type I IFNs. Absence of IFNAR1 exacerbated lysosomal damage in both LPS and poly(I:C)-stimulated BMM, substantially reducing the protective effect of either agonist (Fig 4). Loss of IFNAR1 led to a shift in the distribution toward high values of damage for BMM treated with either LPS or poly(I:C) (Suppl Fig 4C, D). Therefore, the type I IFN response contributes to renitence induced by stimulation of TLRs 4 and 3.

**Figure 4.**
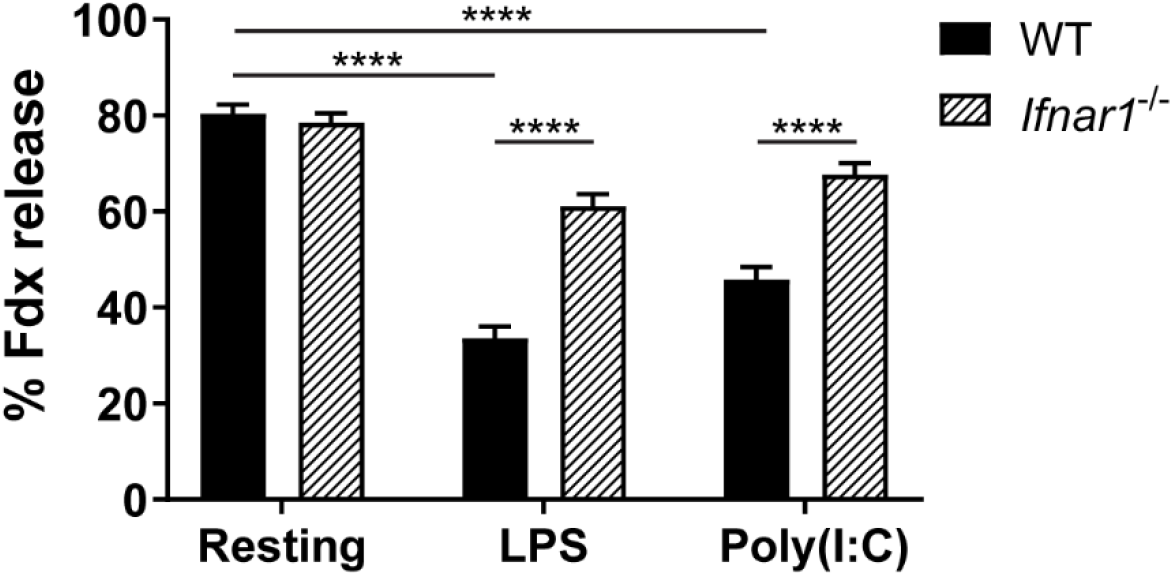
Renitence induced by stimulation of TLR4 or TLR3 depends on the type I IFN response. C57BL/6J (WT) BMM and *Ifnar1*^-/-^ BMM were treated overnight with either LPS (a TLR4 agonist) or poly(I:C) (a TLR3 agonist) or left untreated while subjected to pulse-chase labeling of lysosomes with Fdx. BMM were then incubated with AW beads for 60 min and assayed for lysosomal damage. Bars represent the average percent Fdx release ± SEM per condition for pooled cell data from three independent experiments (n > 252 cells per condition). \*\*\*\**p* ≤ 0.0001.

### MNV-1 infection induces renitence

Poly(I:C) is a synthetic analog of double-stranded RNA (dsRNA), which is carried within viral genomes or represents an intermediate of viral replication of single-stranded RNA (ssRNA) viruses. Its recognition by innate immune receptors (by TLR3 within endosomes, or RIG-1 or MDA5 intracellularly) triggers type I IFN signaling and the induction of anti-viral responses [26]. Previously we discovered that infection of macrophages with hemolysin-deficient *L. monocytogenes*, which cannot perforate phagolysosomes, conferred protection from lysosomal damage upon subsequent challenge with silica beads [7]. Here we investigated whether an analogous protective effect may be conferred by viral infection - that is, whether sublethal exposure to viral infection protects against subsequent lysosomal damage challenge.

To address this question, we assayed lysosomal damage responses in BMM first subjected to viral infection with murine norovirus-1 (MNV-1), which has been shown to infect macrophages and whose replication in macrophages is restricted by type I IFN signaling [27]. BMM were infected with MNV-1 at three different MOIs (0.05, 0.5, 5) for 1 hour, and then were washed and subjected to overnight pulse-chase labeling of lysosomes. Compared to resting BMM, BMM first subjected to MNV-1 infection showed enhanced protection against lysosomal damage at all MOIs tested, although to a lesser degree than that offered by LPS (Fig 5). Renitence capacity increased with increasing viral load, which was confirmed by measurement of viral titers from macrophages infected in parallel as those assayed for lysosomal damage (Suppl Fig 6). As seen for other factors that induce renitence, MNV-1 infection enabled a predominant population of macrophages to preserve their lysosomal integrity following exposure to a lysosomal damaging agent (Suppl Fig 5).

**Figure 5:**
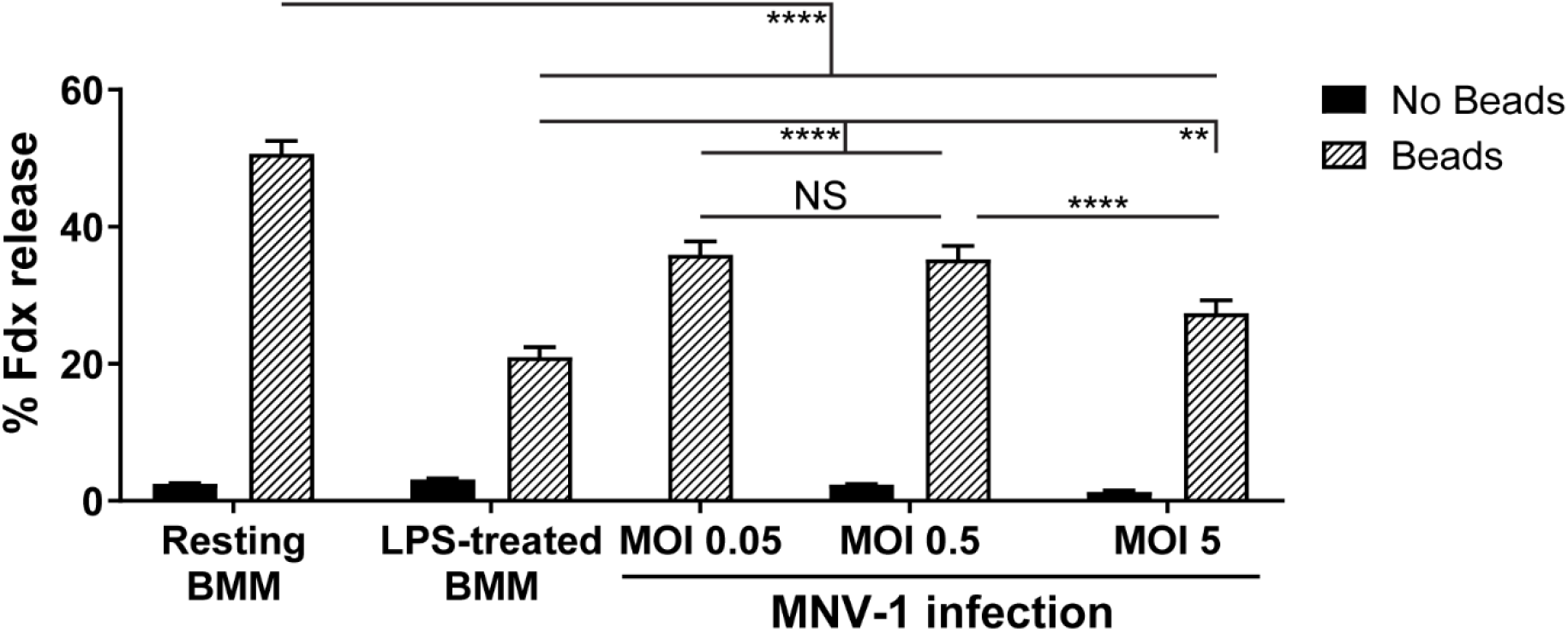
MNV-1 infection induces renitence. BMM were infected with MNV-1 at three different MOIs (0.05, 0.5, and 5) for one hour, washed, and then subjected to pulse-chase labeling of lysosomes with Fdx. BMM were then incubated with AW beads or no beads for 60 min and assayed for lysosomal damage. As controls, lysosomal damage was also measured in resting and LPS-treated BMM. Lysosomal damage for cells not receiving beads was not measured for BMM infected with MOI 0.05. Bars represent the average percent Fdx release ± SEM per condition for pooled cell data from five independent experiments. In bead-positive conditions, cells containing three to seven beads were analyzed (n > 396 cells per condition). \*\**p* ≤ 0.01, \*\*\*\**p* ≤ 0.0001.

## Discussion

By systematically evaluating the inflammatory state and renitence capacity of a large range of activated macrophages, this study refined our understanding of the immunological contexts of renitence. Given the large number of pathogens capable of perforating phagolysosomes, we predicted that macrophages exposed to microbial stimuli or undergoing classical activation would display a high capacity for renitence. Indeed, macrophages stimulated with agonists of TLRs 2/1, 3, and 4, as well as CA-Mφ, exhibited renitence. We also discovered that regulatory macrophages were similarly equipped to induce renitence. These macrophages, known for their immunosuppressive function, were closely related in their activation state to macrophages activated overnight with LPS. Other subtypes examined displayed a lesser capacity to protect their lysosomes from damage. Modest protection was observed in macrophages stimulated with agonists of TLRs 7/8 and 9. AA-Mφ were the least renitent, exhibiting a similar susceptibility to lysosomal damage as that seen in resting macrophages.

Reg-Mφ are immunoregulatory macrophages with key roles in suppressing the immune response following clearance of an infection. Their generation is thought to not occur *de novo* at the onset of immune resolution, but instead to involve the reprogramming of inflammatory macrophages, generated through exposure to TLR stimuli, into immunosuppressive macrophages upon recognition of a second signal that stimulates this transition. Considered in this way, CA-Mφ and Reg-Mφ should not be viewed as separate entities formed under disparate contexts, but as different states that can be assumed by the same macrophage at different stages of the immune response. If this model is correct, these two macrophage subtypes, examined here *in vitro*, could represent predominant cell types at different stages of the immune response *in vivo*. That is, whereas Reg-Mφ likely represent macrophages present at the resolution of an immune response, CA-Mφ likely represent macrophages present at the peak of the immune response, generated after the recruitment of NK cells and Th1 cells, the major sources of IFN-γ *in vivo*, to the site of infection. To complete the model, we propose that macrophages present in the early stages of infection might be represented by macrophages stimulated briefly (eg. for one to two hours) with LPS. Macrophages receiving such short-term treatments with LPS were previously noted for their inability to exhibit renitence [7]. Thus, they represent a different activation state than that embodied by either CA-Mφ or Reg-Mφ.

By mapping our knowledge of the susceptibility to lysosomal damage in each of these three macrophage subtypes to a framework in which these subtypes represent macrophages involved in early, intermediate, and late stages of infection, we can track how renitence capacity within macrophages may vary throughout the course of infection. This temporal framework serves as the foundation of a model we propose for the functional relevance of renitence at different stages of the immune response. This model is summarized in Figure 6.

**Figure 6:**
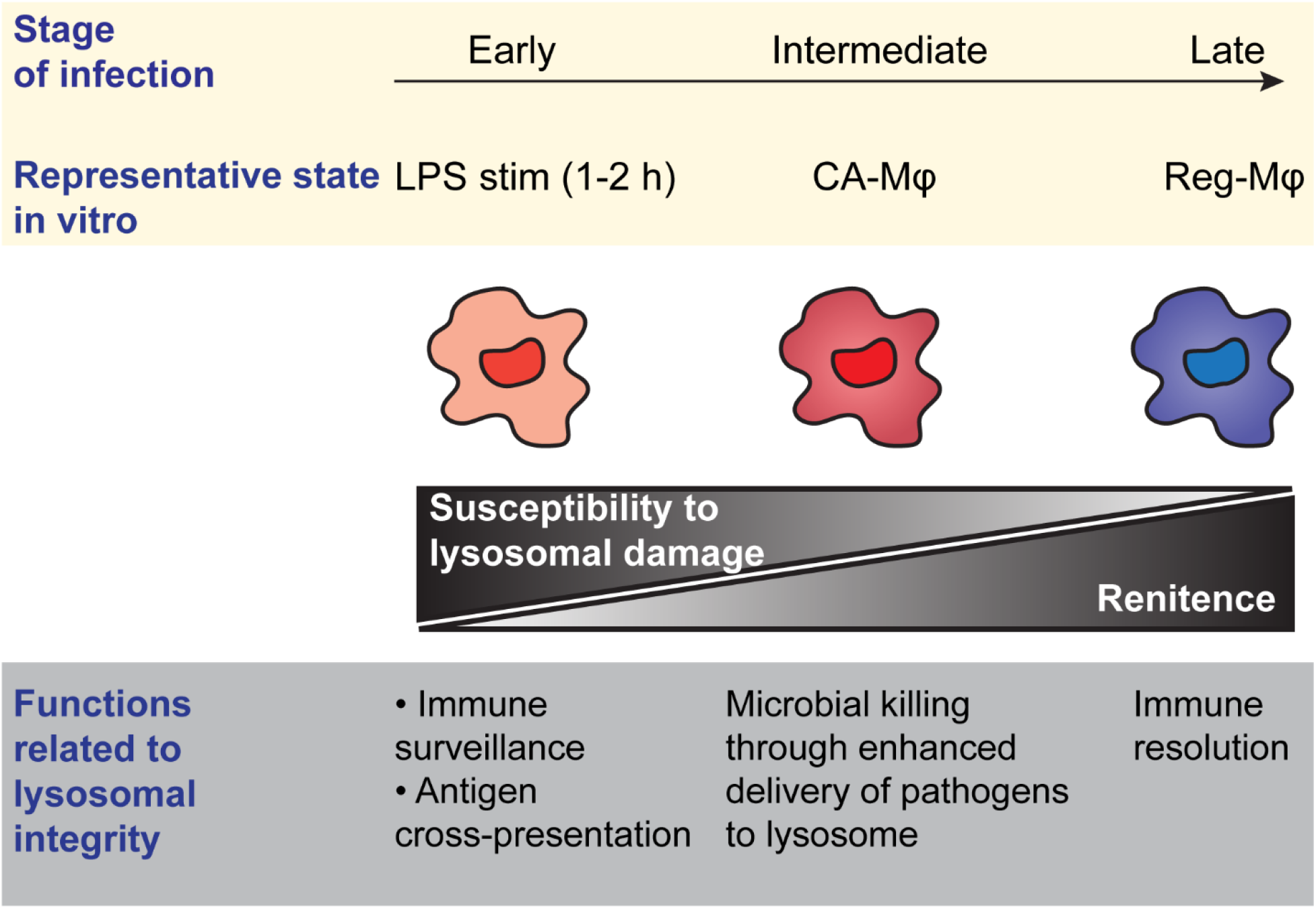
Renitence capacity correlates with macrophage functional state at different stages of infection. Synthesis of our understanding of the macrophage populations examined here and in previous work suggests a model in which three *in vitro* populations of macrophages represent the predominant form of macrophages formed at early, intermediate, and late stages of infection *in vivo*. These, in order, are macrophages stimulated for 1 to 2 h with LPS, classically activated macrophages (CA-Mφ), and regulatory macrophages (Reg-Mφ). In the above model, these macrophages are arranged in their temporal order of generation in the immune response. Overlaying this temporal framework with the renitence capacity for each macrophage subtype allows us to visualize changes in renitence capacity through the course of the immune response. According to this model, renitence is absent early in infection, is upregulated at intermediate stages of infection, and is highest at the stage of immune resolution. In other words, susceptibility to lysosomal damage, considered here as the inverse of renitence, is highest in early stages of infection, and decreases as the infection progresses and resolves. This pattern reflects the immune functions of the predominant macrophage class at each stage of infection, as detailed further in the Discussion section of the text.

Within this temporal framework, renitence is absent in early stages of infection, first observed in macrophages following IFN-γ exposure, and highest in Reg-Mφ involved in the resolution of inflammation. This pattern of lysosomal damage susceptibility, we propose, is consistent with the functional role assumed by macrophages at each stage of infection.

Early during an infection, the main functions of macrophages are the detection of infection and coordination of a proportionate response. A lysosomal network that is more rather than less susceptible to damage may facilitate both functions. For example, the release of microbial products from endolysosomes allows for their detection by inflammasomes in the cytosol, thereby alerting cells of a potential infection by an intracellular pathogen. Although excessive lysosomal damage is deleterious for the cell, trace amounts of leakage that allow for the detection of infection likely promote host defense. The heterogeneous response to damage seen with renitence, in which a subset of cells remains protected and another subset undergoes extreme damage, might enable a coordinated population-level response against an infectious threat. By one model, cells undergoing extreme damage release pro-inflammatory cytokines and danger signals indicating the presence and severity of the threat, thereby amplifying the immune response, whereas the subpopulation of cells undergoing low damage facilitates the prompt restriction of infection and prevention of excess immunopathology.

Over the course of the immune response, the proportion of renitent cells increases, reflecting a higher priority for pathogen restriction and minimizing immunopathology. Pathogen restriction is most effectively accomplished by CA-Mφ. Indeed, exposure to IFN-γ, a necessary signal for CA-Mφ generation, is also necessary for macrophages to restrict infection by several intracellular pathogens. For example, IFN-γ treatment of macrophages infected with *Listeria monocytogenes* prevents escape of the pathogen from phagolysosomes [28]. Likewise, IFN-γ treatment releases the block on phagosome-lysosome fusion imposed by *Mycobacterium tuberculosis* [29, 30]. In both situations, the increased load of pathogen delivery to lysosomes (due to fewer pathogens being lost by escape or stalled in phagosomes) would reasonably be accompanied by an increased capacity by the macrophage to avoid lysosomal membrane perforation by pathogens or their expressed virulence factors.

Following pathogen clearance, the goal of the immune system is to limit rather than to sustain inflammation. Reg-Mφ are key contributors to this process of immune resolution. Shifting the distribution of heterogenous responses to lysosomal damage such that macrophages protected against lysosomal damage make up the predominant sub-population would help to facilitate immune resolution through minimizing possible triggers of inflammation.

The model presented here relies on the assumption that the *in vitro* macrophage subtypes examined indeed represent physiologically-relevant cell types that predominate in a given state of infection. To directly test this model, *ex vivo* analysis of macrophages isolated from *in vivo* mouse models of infection could be pursued. Future work will extend these studies to tissue-specific macrophages as well as to human macrophages in healthy and various disease states.

Our studies of TLR stimulation and renitence we predicted would serve as a basis for interrogating the signaling pathways contributing to renitence. However, the set of renitent vs. non-renitent TLR-stimulated macrophages did not group according to any known parameters associated with TLR signaling. TLRs 2/1, 3, and 4, all of which induce renitence, differ in the signaling adaptors they recruit (TLR2/1 signals through MyD88, TLR3 through TRIF, and TLR4 through either MyD88 at the cell surface or TRIF from within endosomes), in their subcellular localizations (TLR2/1 becomes activated at the cell surface, TLR3 within endosomes, and TLR4 at either location), and in the class of ligands they recognize (TLR2/1 recognizes various ligands found in gram-negative and gram-positive bacteria, mycobacteria and yeast; TLR3 recognizes dsRNA, found in viruses; TLR4 recognizes lipopolysaccharide, a cell wall component of gram-negative bacteria) [22]. Neither do the TLRs whose activation did not profoundly induce renitence share obvious commonalities. TLRs 7/8 and 9 are both endosomally located, but recognize nucleic acids of different classes: TLR 7/8 recognize viral ssRNA as a natural ligand, whereas TLR9 recognizes DNA containing unmethylated CpG motifs, found more abundantly in bacterial and viral than mammalian DNA [22]. The induction of renitence by certain TLR agonists but not others suggests renitence is not a mechanism uniformly upregulated by TLR activation.

The pattern of signaling adaptor requirements revealed here for TLR-induced renitence deviates from the well-established adaptor requirements for TLR signaling leading to NF-κB activation. The main points of deviation lie with the endosomal TLRs 3 and 7/8: the dependence on MyD88 for poly(I:C)-induced renitence, and the requirement for TRIF for R848-induced renitence. These results suggest that cross-talk between MyD88 and TRIF may be necessary for renitence induced downstream of these TLRs. A role for MyD88 in negatively regulating the inflammatory response associated with poly(I:C)-induced TLR3 signaling has been reported in a model of corneal inflammation [31] as well as in immortalized BMM [32]. However, this negative regulation of TRIF signaling by MyD88 was absent in primary mouse BMM [31]. Likewise, the dependence of R848-induced TLR7/8 signaling on TRIF has to our knowledge never been described. The deviation we observe could possibly be explained by a novel pathway of cross-talk between MyD88 and TRIF.

The dependence of TLR4 and TLR3-induced renitence on the type I IFN response suggests renitence may be an innate immune effector function mediated by a set of interferon-stimulated genes (ISGs). Several ISGs have been identified that encode proteins involved in inhibition of endosomal entry of viruses. These include IFITM3, cholesterol 25-hydroxylase, and NCOA7 [33]. Such proteins might also contribute to protection against phagolysosomal damage in settings other than viral infection and may underlie the mechanism of renitence.

The exact mechanism of renitence is unclear but may involve the formation of spacious vacuolar compartments, or “renitence vacuoles,” adjacent to silica bead-containing phagosomes that physically prevent damage or assist in the rapid repair of damaged phagolysosomal membranes [16]. These vacuoles were observed to form and persist in LPS-treated but not resting BMM following uptake of silica beads [16]. Based on analysis of the frequency distributions of damage in protected versus unprotected macrophage subsets, all protected subsets examined seem to restrict damage in a consistent manner – by maintaining a large subpopulation of cells that are protected from lysosomal damage. The mechanism promoting resistance to damage within these protected subsets is unknown but may relate to their ability to induce the formation of renitence vacuoles. Furthermore, how and why renitence induction produces a heterogeneous, as opposed to homogeneous, response to lysosomal damage is unclear. That is, why aren’t 100% of cells within a renitent population protected from damage? Heterogeneous cytokine responses to LPS stimulation in phenotypically homogeneous populations of murine BMM have previously been described [34, 35]. In one study, IL-12p40 production in LPS-stimulated BMM at the population level was shown to assume a bimodal distribution, with the exact proportion of high versus low IL-12p40-producing cells controlled by the phase of the intracellular molecular clock [34]. Another recent study, in which macrophage activation was assessed by expression of TNF or RelA (the p65 subunit of NF-κB), reported a bimodal distribution of responses to LPS stimulation that is regulated by cell density [35]. Whether the readout is cytokine secretion or lysosomal damage protection, the ability of innate immune cells to mount a heterogeneous response to microbial infection at the population level likely enables a more fine-tuned and adaptable immune response. As proposed for our working model above, in the case of renitence, it is conceivable that the subset of cells undergoing damage serves to provide signals alerting the immune system, and the remaining undamaged cells, of the presence, nature, and severity of infection.

The observation that MNV-1 infection protects against subsequent phagolysosomal damage in macrophages supports the concept that sublethal viral infection can prime cells to defend against future membrane-damaging threats. Whether the induction of renitence promotes the restriction of viral escape from endosomes is not known, and can be investigated using established models for measuring the extent of MNV-1 endosomal escape [36].

Distinct mechanisms for inducing renitence may exist and vary depending on the microbial stimulus sensed and innate immune pathway activated. The pattern recognition receptor that recognizes MNV-1 is a question of active investigation. MNV-1 is a ssRNA virus whose recognition is mediated by the intracellular sensor MDA5, which traditionally recognizes dsRNA [37]. As such, MDA5 presumably recognizes a replication intermediate of MNV-1. Whether MNV-1 has a cognate Toll-like receptor and whether stimulation of intracellular sensors induces renitence are unknown. It is conceivable that stimulation of cytosolic sensors (e.g. by MNV-1) might promote renitence through a different mechanistic pathway than stimulation of TLRs (e.g. by LPS).

In summary, this study characterized the range of macrophage populations capable of undergoing renitence. Consideration of how the populations examined correspond to macrophages at different temporal stages of the immune response suggests a model in which renitence capacity increases as an infection progresses and resolves. Together, this work highlights the significance of a poorly understood cell biological property – susceptibility to lysosomal damage – and positions us to continue to uncover how this property reflects and contributes to the diverse functional roles of macrophages.

## Authorship

A.O.W. participated in the conceptualization, investigation, data analysis, and original writing of the manuscript as well as subsequent revisions. M.M. and I.A.O. participated in the investigation, data analysis, and revisions of the manuscript. C.E.W. and J.A.S. participated in the conceptualization and data analysis, as well as the review, editing and revision of the manuscript.

## Acknowledgments

The authors thank David Friedman, Holly Turula, Zachary Mendel, and Michele Swanson for helpful discussions. This work was supported by NIH grants R01-GM101189 and R35-GM131720 to J.A.S., and the Biological Sciences Scholars Program at the University of Michigan to C.E.W. A.O.W. was supported by the Mechanisms of Microbial Pathogenesis training program (T32-AI-007528) and the University of Michigan Rackham Predoctoral Fellowship program. I.A.O. was funded by a Michigan Infectious Disease International Scholars fellowship from the Department of Microbiology and Immunology at the University of Michigan and a DELTAS Africa grant (DEL-15-007: Awandare) under the Wellcome Trust to the University of Ghana.

