## Supplemental figures for "Macrophage Inflammatory State Influences Susceptibility to Lysosomal Damage"

### Supplemental Materials
***Journal of Leukocyte Biology***

Wong et al.

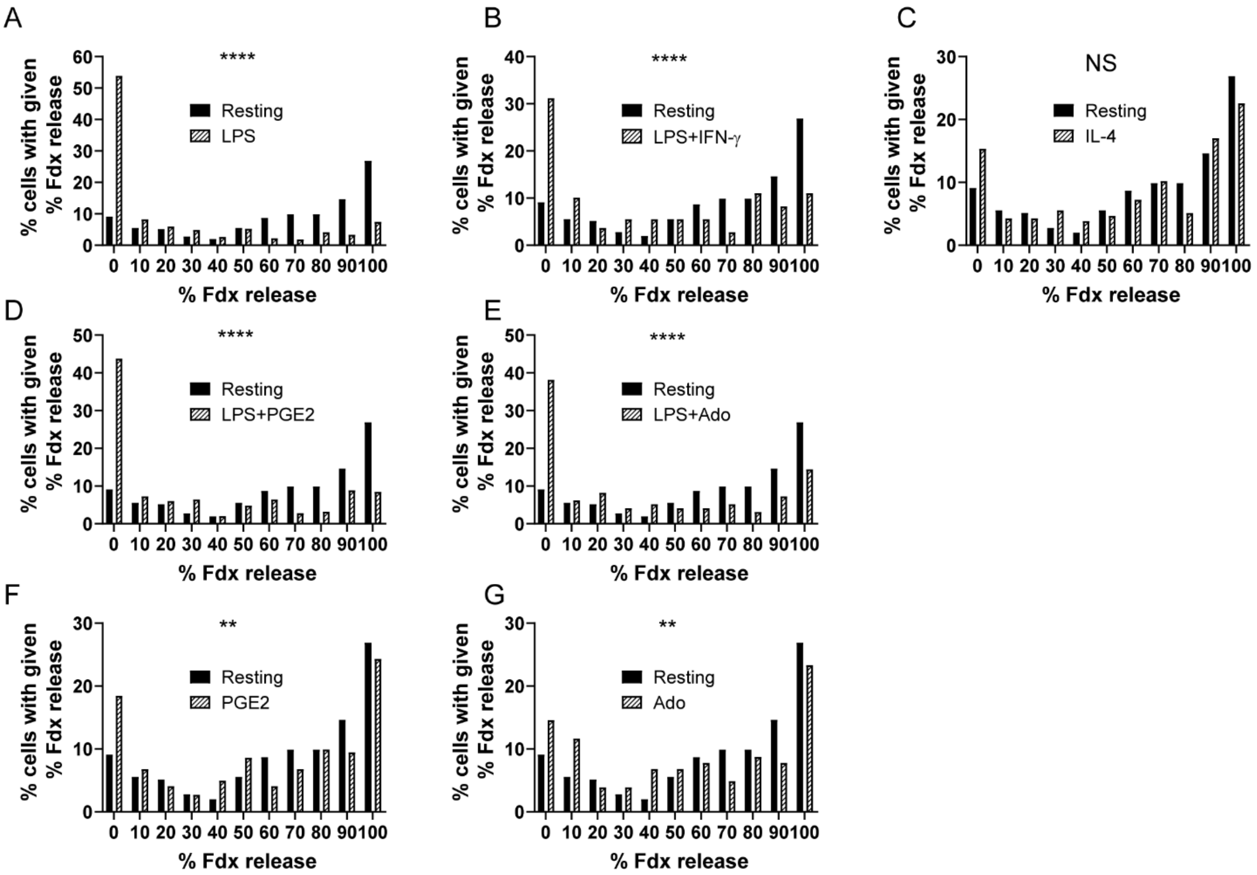


**Supplemental Figure 1: Frequency distributions of data sets in Figure 1C.**

For each bead-positive condition analyzed in Figure 1C, individual cell data was plotted in histogram form to depict the percent of cells within each condition experiencing a given level of lysosomal damage. The relative frequency in percentages (y-axis) of a given level of % Fdx release (represented in increments of 10% on the x-axis) for each treatment condition was plotted and compared with the frequency distribution of % Fdx release values for resting BMM. The frequency distribution for resting BMM was compared with that for BMM stimulated with LPS (A), LPS+IFN-γ (B), IL-4 (C), LPS+PGE2 (D), LPS+Ado (E), PGE2 (F), Ado (G). Statistical significance of the difference between each pair of frequency distributions was determined using the Kolmogorov-Smirnov test. Shown are pooled cell data from two to four independent experiments (n > 97 cells per condition). NS: no significant difference, $**p \leq 0.01$, $****p \leq 0.0001$.

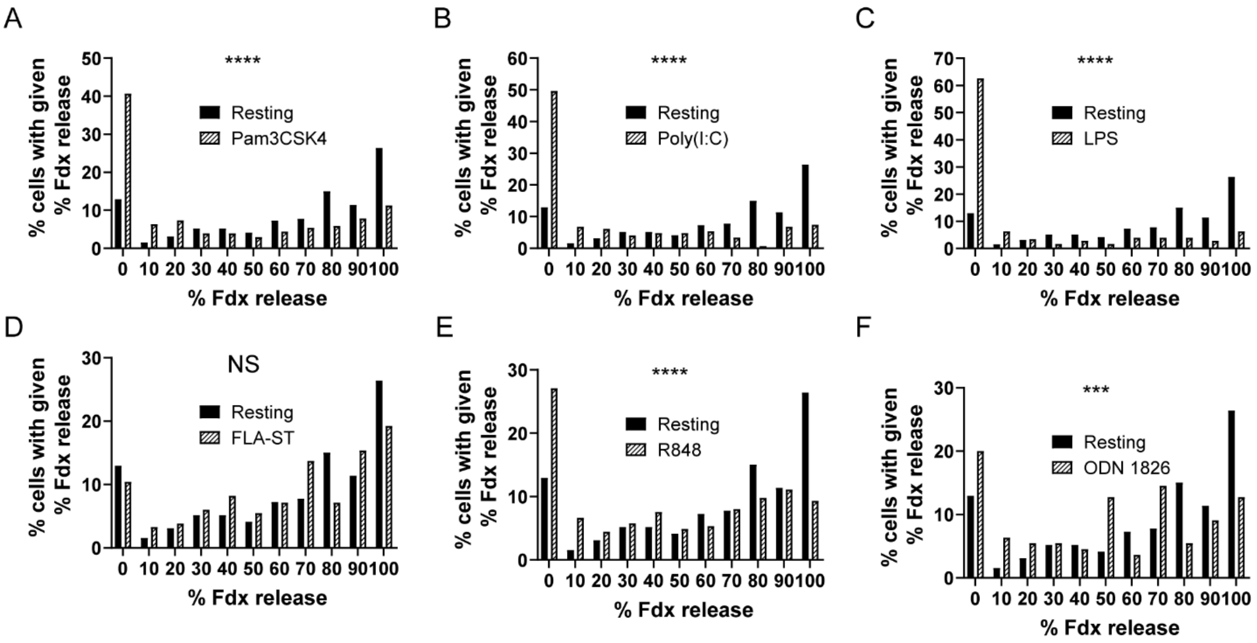


**Supplemental Figure 2: Frequency distributions of data sets in Figure 2A.**

Histograms depicting the percent of cells (y-axis) experiencing a given level of % Fdx release (represented in bins of 10% on the x-axis) based on individual cell data from each bead-positive condition analyzed in Figure 2A. The frequency distribution for resting BMM was compared with that for BMM stimulated with the following TLR agonists: (A) Pam3CSK4, (B) Poly(I:C), (C) LPS, (D) FLA-ST, (E) R848, (F) ODN 1826. Shown are pooled cell data from two or three independent experiments (n > 110 cells per condition). NS: no significant difference, ***$p \leq 0.001$, ****$p \leq 0.0001$.

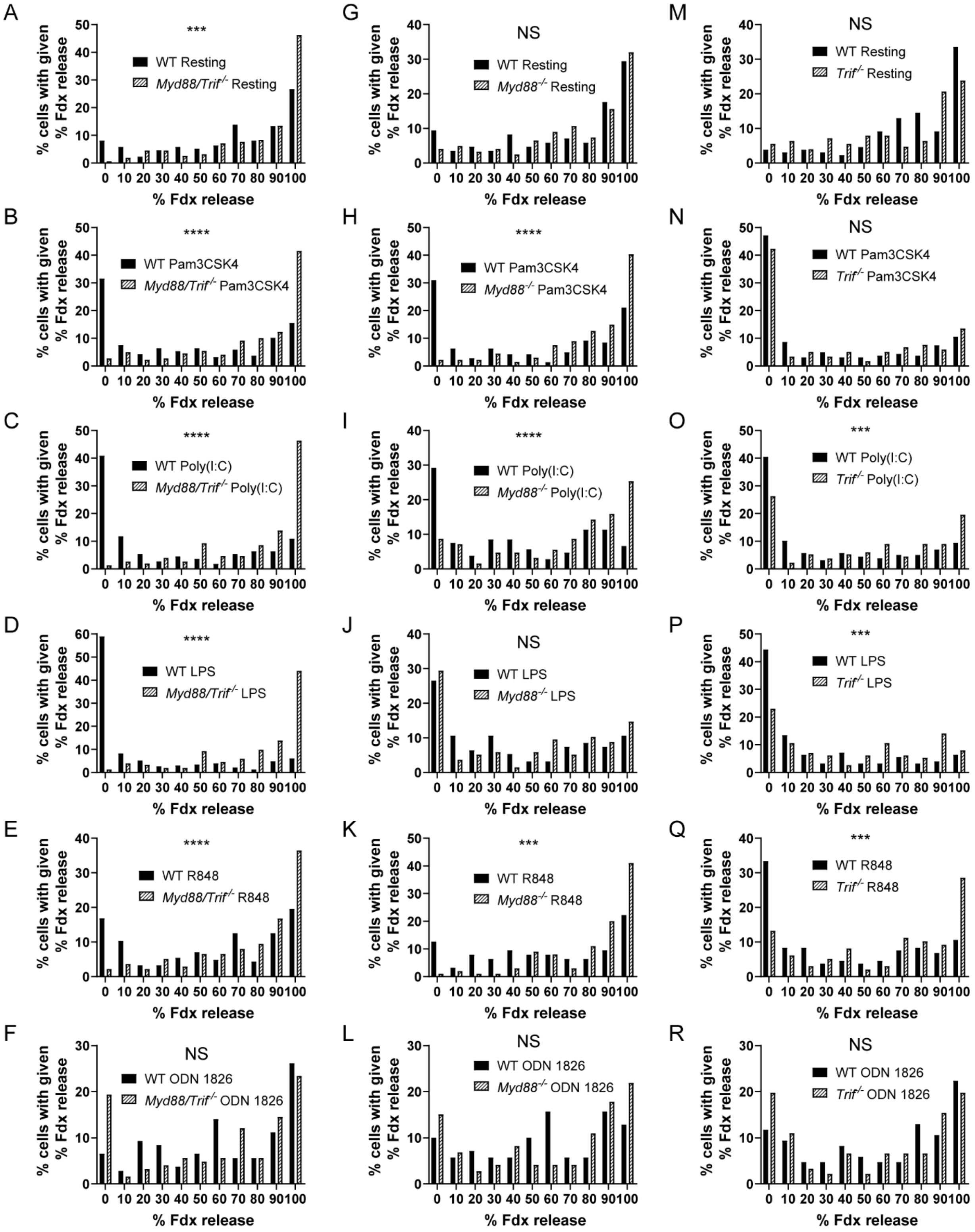
A
***
WT Resting
Myd88/Trif-/- Resting
% cells with given % Fdx release
% Fdx release
G
NS
WT Resting
Myd88-/- Resting
M
NS
WT Resting
Trif-/- Resting
B
****
WT Pam3CSK4
Myd88/Trif-/- Pam3CSK4
H
****
WT Pam3CSK4
Myd88-/- Pam3CSK4
N
NS
WT Pam3CSK4
Trif-/- Pam3CSK4
C
****
WT Poly(I:C)
Myd88/Trif-/- Poly(I:C)
I
****
WT Poly(I:C)
Myd88-/- Poly(I:C)
O
***
WT Poly(I:C)
Trif-/- Poly(I:C)
D
****
WT LPS
Myd88/Trif-/- LPS
J
NS
WT LPS
Myd88-/- LPS
P
***
WT LPS
Trif-/- LPS
E
****
WT R848
Myd88/Trif-/- R848
K
***
WT R848
Myd88-/- R848
Q
***
WT R848
Trif-/- R848
F
NS
WT ODN 1826
Myd88/Trif-/- ODN 1826
L
NS
WT ODN 1826
Myd88-/- ODN 1826
R
NS
WT ODN 1826
Trif-/- ODN 1826

**Supplemental Figure 3: Frequency distributions of data sets in Figure 3.**

Histograms depicting the percent of cells (y-axis) experiencing a given level of % Fdx release (represented in bins of 10% on the x-axis) based on individual cell data from each bead-positive condition analyzed in Figure 3. To visualize the effect of each genetic knockout on the distribution of damage for each stimulation condition, the frequency distribution for a given stimulation condition was compared between WT and *Myd88/Trif*$^{-/-}$ BMM (A-F), WT and *Myd88*$^{-/-}$ BMM (G-L), and WT and *Trif*$^{-/-}$ BMM (M-R). Shown are pooled cell data from two independent experiments (n > 107 cells for A-F, n > 63 cells for G-L, n > 85 cells for M-R). NS: no significant difference, ***$p \leq 0.001$, ****$p \leq 0.0001$.

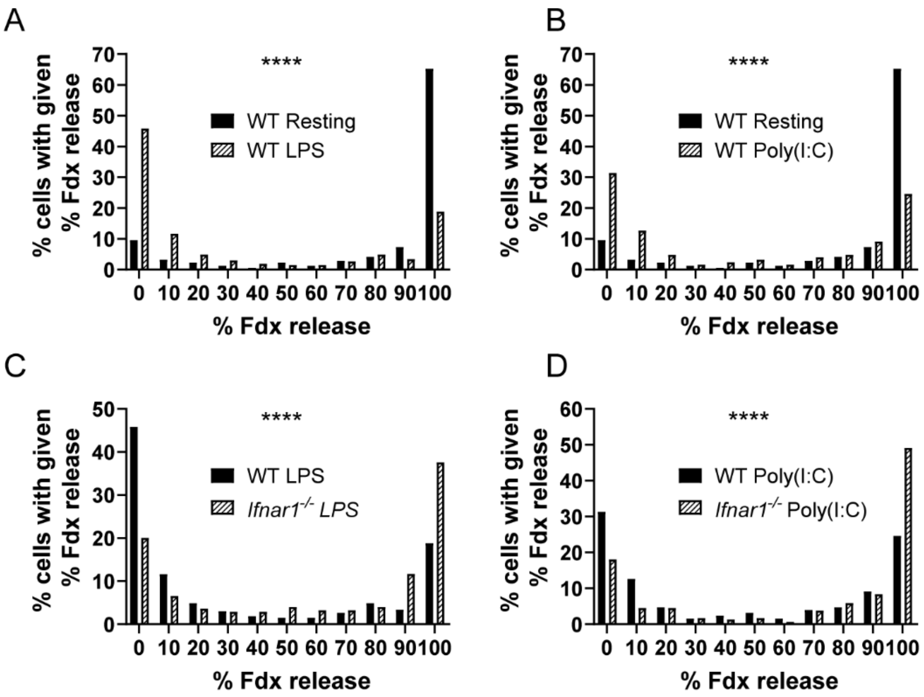


**Supplemental Figure 4: Frequency distributions of data sets in Figure 4.**

Histograms depicting the percent of cells (y-axis) experiencing a given level of % Fdx release (represented in bins of 10% on the x-axis) based on individual cell data from each bead-positive condition analyzed in Figure 4. In the top panel, the frequency distribution for resting WT BMM was compared with that for WT BMM stimulated with LPS (A) or poly(I:C) (B). In the bottom panel, the frequency distribution for each given stimulation condition was compared between WT and *Ifnar1*$^{-/-}$ BMM. Shown are pooled cell data from three independent experiments (n > 252 cells per condition). ****$p \leq 0.0001$.

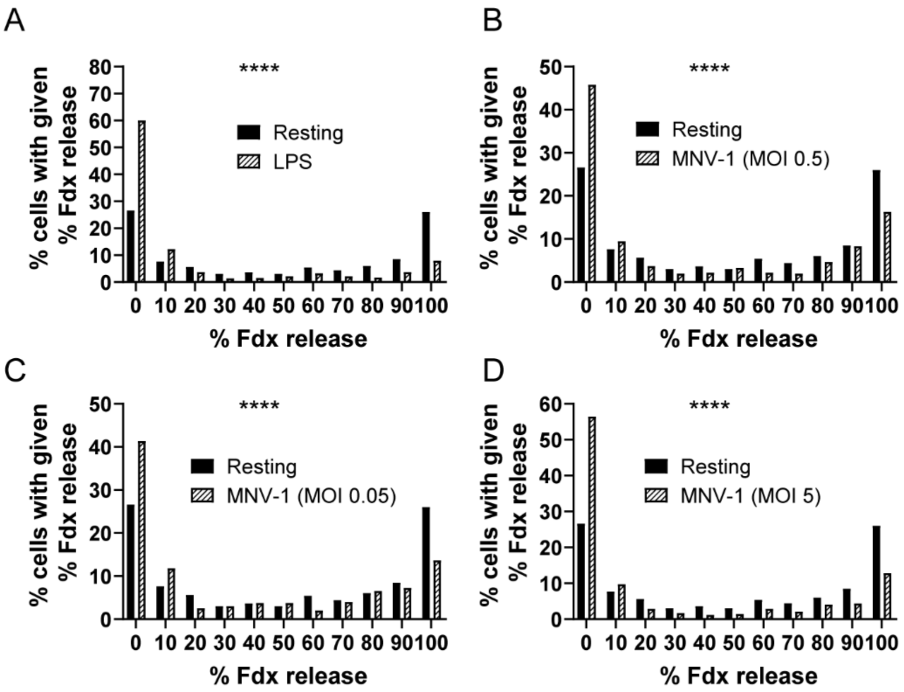


**Supplemental Figure 5: Frequency distributions of data sets in Figure 5.**

Histograms depicting the percent of cells (y-axis) experiencing a given level of % Fdx release (represented in bins of 10% on the x-axis) based on individual cell data from each bead-positive condition analyzed in Figure 5. The frequency distribution for resting BMM was compared with that for BMM stimulated with LPS (A) or infected with MNV-1 at one of three MOIs (0.05, 0.5, 5; B-D) before being subjected to lysosomal damage. Shown are pooled cell data from five independent experiments (n > 396 cells per condition). ****$p \leq 0.0001$.

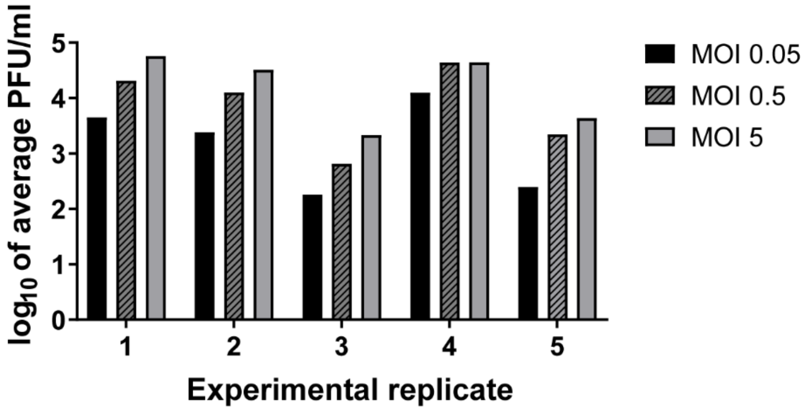


**Supplemental Figure 6: MNV-1 infects BMM in an MOI-dependent manner at 18 hours post-infection in 5 different experiments.**

WT BMM were infected with MNV-1 at three different MOIs (0.05, 0.5 and 5). Viral titers in cell culture lysates were measured by virus titration using a plaque assay and reported as plaque forming units/ml (PFU/ml). Bars show MNV-1 infection titers of three different MOIs from 5 independent experiments performed in duplicate or triplicate. These assays were performed in parallel with the viral infections for the lysosomal damage experiments described in Figure 5.
